# Real-time visualisation of the intracellular dynamics of *Shigella’s* virulence plasmid

**DOI:** 10.1101/2025.07.10.664258

**Authors:** Tanuka Sen, Valentine Lagage, Stephan Uphoff, Christoph Tang

## Abstract

Bacterial plasmids play critical roles in horizontal gene transfer, antibiotic resistance, virulence, and in shaping the interaction of bacteria with their environment, including those encountered in their hosts. They are integral to bacterial evolution as they act as vectors for rapid genetic change and adaptation. *Shigella sonnei* is a human-adapted pathogen that is a leading cause of bacillary dysentery and a significant threat to public health. In common to all species of *Shigella*, *S. sonnei* harbours an ∼ 210 kb virulence plasmid, pINV, which is essential for its virulence. pINV encodes a Type III secretion system that mediates bacterial invasion of epithelial cells. Understanding the dynamics of replication and segregation of pINV is therefore necessary to define how *Shigella* retains the plasmid and thence virulence. In this study, we used three fluorescent tagging approaches to detect the presence of pINV in live *S*. *sonnei* to allow tracking of the plasmid during cell division. Conclusively, we found that tagging the plasmid with the *parS-*ParB partitioning system sequences from *Caulobacter crescentus* chromosome, allowed suitable visualization of the single copy pINV in *S. sonnei.* This comprehensively enabled us to monitor the location and segregation of the plasmid during bacterial growth.

## INTRODUCTION

*Shigella* is a Gram-negative, human-adapted bacterial pathogen, that causes a spectrum of symptoms ranging from mild gastroenteritis to severe dysentery [1]. There are four known species of *Shigella*, with *Shigella flexneri* and *Shigella sonnei* accounting for the majority cases worldwide [1, 2]. Overall, *Shigella* is a leading contributor to the global burden of diarrhoeal infection in children [3, 4]. Every year, *Shigella* causes 80–165 million cases of shigellosis and between 6,900–30,000 deaths worldwide [2, 3]. A fundamental player in the virulence of *Shigella* is a 210-230 kb virulence plasmid, pINV [4]. This genetic element orchestrates an array of molecular events that enable *Shigella* to invade into epithelial cells, spread from cell-to-cell, then manipulate the host cytoskeleton and inflammatory responses [4, 5].

*Shigella* is known to evolve from *Escherichia coli* following acquisition of pINV which encodes for a range of factors needed for host cell invasion and the intracellular lifestyle of *Shigella* [5-7]. A conserved 31 kb pathogenicity island (PAI) on pINV, contains the *ipa-mxi-spa* loci, encoding a Type 3 Secretion System (T3SS) and several protein effectors delivered through the T3SS [7-9]. Through the multifaceted actions of pINV-borne virulence genes, *Shigella* gains access to an intracellular niche, subverts host defences, induces pyroptosis in macrophages, and initiates a cascade of events that lead to clinical disease [7-9]. The copy number of pINV is strictly controlled at 1-2 copies per cell, as expression of the T3SS and its related genes impose a metabolic fitness cost on bacteria; loss of pINV or the T3SS PAI leads to enhanced growth of bacteria [9, 10]. *S. flexneri* is known to have at least five plasmid maintenance systems, which include three type II toxin–antitoxin (TA) systems, *vapBC* (also known as *mvpAT*), *ccdAB,* and *gmvAT*, as well as two partitioning systems *parAB* and *stbAB* [10, 11]. *S. sonnei* in comparison employs one or two TA systems (*vapBC* and *relBE*) and one partitioning system (*parAB*) that maintain pINV in a population of dividing bacteria [10, 11]. The TA systems operate as “plasmid addiction” elements that kill or inhibit the growth of bacteria that lose the plasmid during cell division, a process known as post-segregational killing (PSK) [10, 11].

Investigating molecular dynamics of a plasmid in real-time is crucial for understanding gene transfer, antibiotic resistance, and the evolution and maintenance of bacterial virulence [12, 13]. Development of cutting-edge imaging technologies has enabled real-time analysis of plasmids in living bacterial cells during cell division and transfer [12, 13]. Given the pivotal role of pINV for *Shigella* during host:pathogen interactions, it is crucial to understand the dynamic interactions between this plasmid and bacteria. In this study, we aimed to examine the replication and spatial distribution of pINV in live *Shigella* at a single-molecule level. To achieve this, we evaluated three approaches to detect the presence of the plasmid using fluorescent tagging in *S. sonnei* CS14. In comparison to other *S. sonnei,* these clinical isolate harbours a stable form of pINV due to a single nucleotide polymorphism in the *vapBC* TA system. This mutation ensures that pINV is efficiently maintained in bacterial populations through highly efficient PSK system. The major driving force of PSK is the antitoxin systems (VapBC) that eliminate cells lacking pINV after division [14]. Each of three tagging systems tested here had their own strengths and limitations. In the present study, we identified a tagging system, which can accurately track the maintenance, loss, location, organisation, and segregation of a low copy number genetic element such as the pINV during bacterial growth.

## MATERIALS AND METHODS

### Bacterial strains and growth

The bacterial strains and plasmids used in this study are shown in Supplementary Table 1. *Shigella* and *E. coli* were grown in liquid Luria Bertani (LB) broth or on solid media containing 1.5% (w/v) agar (Oxoid, Basingstoke, UK). For detecting expression of the T3SS and plasmid loss assays, *Shigella* was grown in tryptic soy broth (TSB), containing 1% Congo red (CR) added to solid media [10, 11]. To avoid auto-fluorescence of LB media, for fluorescence microscopy bacteria were grown in glucose minimal media (GMM) supplemented with essential amino acids: methionine 45 μg/mL, tryptophan 125 μg/mL, and nicotinic acid 20 μg/mL. Bacterial cultures were incubated at 37°C, unless bacteria contained a temperature sensitive plasmid (pKD46), which was only grown at 30°C. Antibiotics were added at the following concentrations: 100 μg/mL ampicillin; 50 μg/mL kanamycin; 25[μg/mL chloramphenicol. For inducing protein expressions, 20% arabinose w/v and 1M IPTG w/v stock solutions were prepared.

### Strain construction

DNA tag sequences were inserted into *S. sonnei* pINV using the phage λ-red homologous recombination system, utilizing the plasmids pKD46 and pKD3 chloramphenicol cassette as the resistance marker [15]. In all the cases, sequences were introduced into pINV at an 8 bp intragenic region between *vapC* and an inactive *traD* gene, which are orientated in a tail-to-tail fashion. Sequences were inserted 4 bp downstream of the *vapC* stop codon which does not affect the function of the TA system [16]. This location specific insertion strategy along with introducing an antibiotic cassette adjacent to a fluorescent tag, ensures the presence of the inserted tag, as it can be selected for during bacterial growth over multiple generations [16]. Constructs were assembled using Gibson Assembly (NEB) and verified by colony PCR and Sanger sequencing (Source BioScience). All primers used in this study are listed in Supplementary Table 2.

### Labelling pINV with a fast-degrading SsrA tag

pROD50 expressing the mYPet fluorescent protein [17] was modified by introducing a ssrA protein degradation sequence. An 11 amino acid peptide (AANDENYALAA) sequence was added to the C-terminus of mYPet [18-20]. This modified pROD50 was used as template to amplify the sequence encoding the mYPet-ssrA fusion, along with a kanamycin resistance cassette via PCR. The primers used are listed in Supplementary Table 2. The amplified 2396 bp sequence was inserted into pINV at the chosen location (Figure 1A). Expression of mYPet-ssrA was induced by adding 0.1 mM IPTG to *S. sonnei* containing pINV encoding the mYPet-ssrA fusion tag at OD600 = 0.2, for 4 hours at 37 °C. A 1mL aliquot of the culture was pelleted, washed, and re-suspend in 20 μL GMM for imaging on agarose pads.

**Figure 1:**
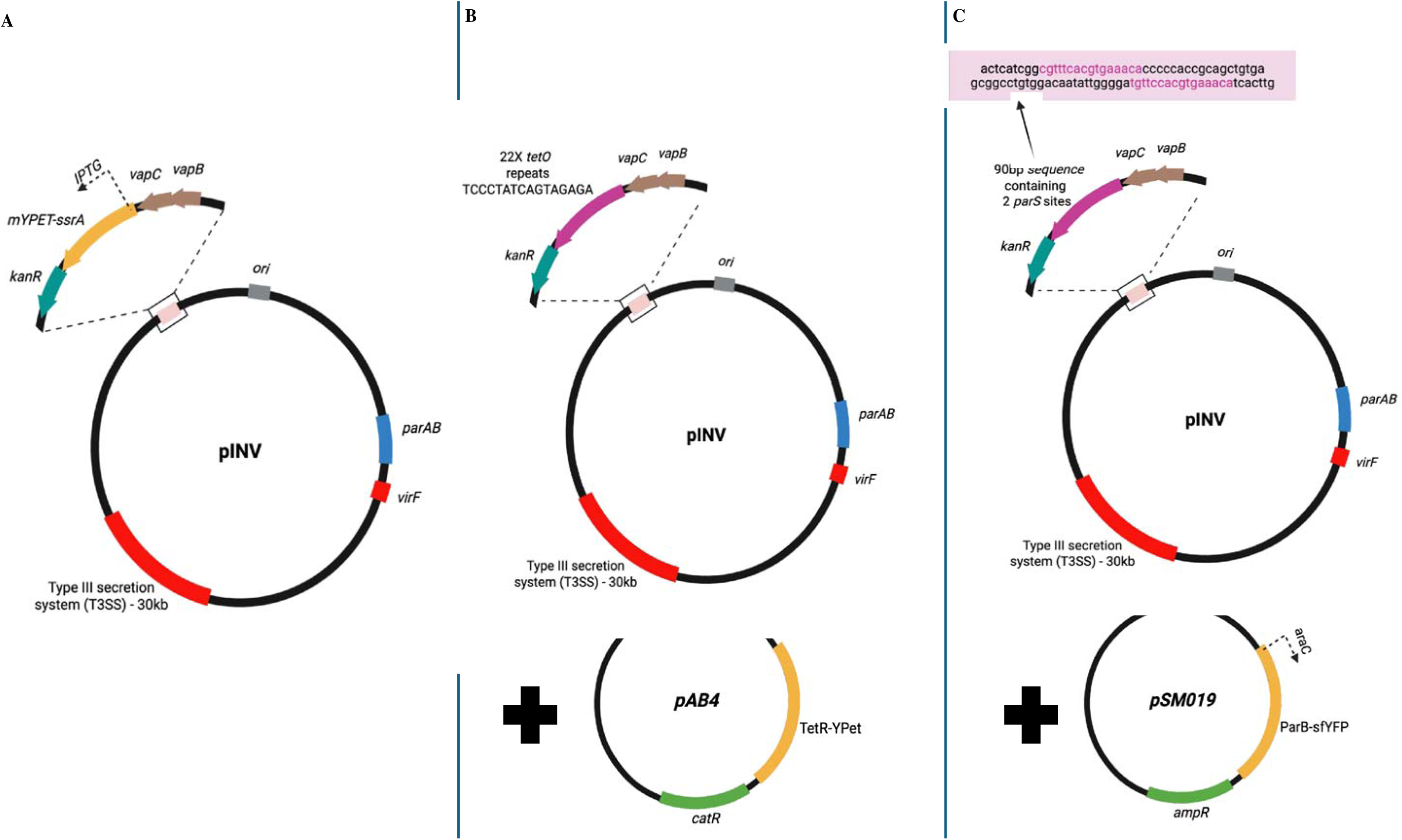
Schematic representation of pINV fluorescent tagging strategies. Construction and configuration of three different pINV tagging systems used for live-cell visualization of *Shigella* cells harbouring the virulence plasmid (pINV). (**A)** mYPet-ssrA tagging system: A C-terminal fusion of the mYPet fluorescent protein with an 11-amino-acid ssrA degradation tag (mYPet-ssrA) was inserted into pINV under the control of an IPTG-inducible promoter. This system enables transient fluorescent labeling of pINV-containing *Shigella* cells, which appear as uniformly fluorescent yellow cells under the microscope. **(B)** *tetO*/TetR-YPet tagging system: A 22x *tetO* operator array was inserted downstream of the *vapBC* locus on pINV. A second plasmid, pAB4, encodes a TetR-YPet fusion protein that specifically binds the *tetO* array, resulting in discrete fluorescent foci (typically 1–2 per cell) corresponding to tagged pINV copies. **(C)** *parS*/ParB-sfYFP tagging system: A 90 bp synthetic sequence containing two *parS* sites was inserted into pINV, serving as a binding site for the ParB-sfYFP fusion protein. The fusion is encoded on the arabinose-inducible plasmid pSM019. Binding of ParB-sfYFP to the *parS* sites results in visible fluorescent foci marking the pINV plasmid within live cells. In all three systems, insertions on pINV were made downstream of the *vapBC* stop codon to preserve the integrity of the native toxin-antitoxin system. Gene and feature annotations: *ori*: origin of replication; *vapBC*: toxin-antitoxin system; *parAB:* plasmid partitioning system; *virF:* virulence regulator. Plasmid resistance markers: *kanR* – kanamycin resistance, *catR* – chloramphenicol resistance, *ampR* – ampicillin resistance.

### Labelling pINV with the *tetO*-TetR system

*E. coli* AB1157 containing pLAU44 [21] was used as the template to amplify 22 *tetO* repeats (each 19 bp) by PCR primers (Supplementary Table 2). The 865 bp *tetO* fragment along with a chloramphenicol resistance cassette were inserted into pINV as above, generating pINV-*tetO*. pAB4 expressing the TetR-YPet fusion protein [21] was then transferred into *S. sonnei* containing pINV-*tetO* (Figure 1B). A 5 mL aliquot of the log phase bacterial culture was pelleted down, washed, and re-suspend in 200 μL of GMM to a final concentration of OD_600_ = 0.2 for imaging on agarose pads.

### Labelling pINV with the *parS*-ParB system

Plasmid pSM040 (*Cm*) [22] containing the *C. crescentus parS* sequence was used as the template in this case. A 90 bp region comprising of two *parS* sites, along with a chloramphenicol resistance cassette was amplified from pSM040 via PCR using specific primers (Supplementary Table 2). This fragment was then inserted into pINV. pSM019 (*amp*^R^) encoding ParB-sfYFP fusion protein [22], under the control of an arabinose inducible promoter was then transformed by electroporation into *S. sonnei* containing the pINV with the *parS* sites (Figure 1C). At OD600= 0.2, the ParB-sfYFP fusion was expressed by inducing with 0.2% arabinose for 2 hours at 37 °C. A 1ml aliquot of OD600 = 0.6 culture was then pelleted, washed, and re-suspended in 200 µl GMM for imaging on agarose pads.

### Live-cell imaging on agarose pads and analysis

Bacteria were added to GMM-1% agarose-mounted slides for live microscopy. A 5 μL aliquot of bacterial culture was spotted onto the agarose pad, left to dry for 5 minutes, after which a coverslip was placed on top of the sample. Images were acquired using an EVOS® FL Auto Imaging System (100X (NA1.4, oil-immersion type lens). The thermostatic microscope chamber was preheated to 37^0^C, and imaging was carried out at this temperature. Fluorescence images were acquired at excitation wavelength of 470/22 nm and emission wavelength of 525/50 nm for YFP based plasmid tags [23].

Images of live bacteria were analysed by ImageJ/Fiji [24] the open-source MicrobeJ plug-in [25] and BacStalk [26]. Segmentation which involved defining the bacterial perimeter; quantitative analysis of the morphology; the fluorescence intensity of cells, and its distribution; and automated detection of fluorescent foci were carried out with these tools. Phase contrast images were used to segment bacterial cells and the fluorescence images capturing pINV with the *parS*-ParB-sfYFP tag were used to detect the presence/absence of pINV, BacStalk was then used for data plotting, creating a demograph representing the movement of pINV across the cell, during cell division.

### Growth and plasmid loss assays

*S. sonnei* was grown overnight in liquid cultures then diluted to 1:50 into 25 mL of fresh LB or GMM, then incubated at 37°C with shaking (180 rpm). Every 60 minutes, the optical density (OD) of the bacterial culture was measured at 600 nm (OD_600_).

To assess the maintenance of pINV in bacteria, plasmid loss assays were performed. *S. sonnei* expressing an active T3SS borne on pINV, bind to CR when grown on solid media containing this dye [10, 11]. *S. sonnei* strains were grown at 37°C on CR-TSB agar plates containing chloramphenicol overnight to obtain single colonies. Three independent CR^+^ colonies were re-suspended in a 5 mL TSB liquid media and incubated at 37°C with shaking at 180 rpm for 25 generations (16 hours). Samples were diluted in PBS and plated onto CR-TSA and incubated overnight at 37°C before counting the number of CR^+^ and CR^−^ colonies.

The proportion of CR^−^ colonies was quantified by dividing the number of emerging CR^−^ colonies by the total number of colonies (CR^+^ and CR^−^) and expressed as a percentage. Colonies were evaluated by visual examination [10, 11]. Furthermore, multiplex PCR was performed with primers designed for pINV genes [14] (Supplementary Table 2) *virF, virB, mvpAT,* and *ori* to generate products of distinct sizes; *hns*, a chromosomal gene, was included as a control [10, 11].

## RESULTS

Our aim was to evaluate methods that could detect the presence of pINV in *S. sonnei* either by labelling cells containing pINV or directly labelling the pINV. Our emphasis was to develop a fluorescent tag that would reliably identify cells that had lost the plasmid, soon after it occurs. This could enable us to track subsequent phenotypic changes in bacterial morphology, growth and metabolism during the process of PSK after plasmid loss. Additionally, the ability to track the plasmid location and copy number of pINV inside bacteria, as well as its movement during and after cell division.

The first tagging approach employed expression of a pINV-encoded fluorescent reporter protein to detect the presence of the plasmid in the cell. The rationale for this approach was that only bacteria harbouring the plasmid will express the fluorescent protein and plasmid loss could be detected via the loss of expression. As fluorescent proteins can have an extended half-life (*e.g.* the half-life of GFP in bacteria is 26 hours [23]), we added a fast-degrading ssrA tag to the C-terminus of a fluorescent protein [18-20]. The ssrA tag is an 11-residue peptide (AANDENYALAA), which is recognised by cellular proteases including ClpXP/ClpAP [18-20]. However, this reporter does not provide information on the localisation or copy number of the plasmid. The other tagging approaches we employed are based on adding specific DNA sequences into a target molecule such as a plasmid or chromosome, which are specifically recognised and bound by a fluorescent protein [27, 28]. This approach allows selective labelling and tracking of individual plasmids [27, 28]. The two systems explored in this study include the Tet repressor protein (TetR) with its cognate *tetO* operator sequence, and the partitioning protein (ParB) binding to *cis*-acting *parS* DNA sequence [21, 20]. To prevent the loss of target DNA sequences (*parS* and *tetO*) from pINV via insertion sequences, it was always added near pINV’s origin of replication (ori), and downstream of the vapBC (TA) genes [16]. Additionally, an antibiotic cassette added nearby the *parS/tetO* sites ensured that these inserted sequences are stably maintained on pINV.

### The mYPet-ssrA fusion protein demonstrates inconsistent fluorescence intensities

The fast-maturing fluorescent protein mYPet-ssrA [17-20] along with a kanamycin resistance cassette was inserted into pINV, immediately downstream of *vapBC*. The expression of this fusion tag was then induced with 0.1 mM IPTG for a duration of 4 hours. The bacteria exhibited yellow fluorescence throughout the cytoplasm, indicating protein expression and the presence of pINV (Figure 2A). To mimic plasmid loss, the inducer IPTG was removed by washing bacteria three times with phosphate buffer saline (PBS) solution, then imaging live cells for the next 2 hours. In principle, loss of pINV should be followed by the rapid degradation of mYPet-ssrA fusion protein and decrease in fluorescence intensity. It was observed that the overall cellular YFP intensity decreased over time (Figure 3A), but the fluorescence did not fall below 50% of its original value over the two hours of imaging. Based on the rate of degradation obtained from this experiment (Fig 3A), the half-life of mYPet-ssrA degradation could be estimated to 164 minutes or 2.7 hours approximately. This decrease was not as rapid as described previously in studies using the ssrA tag with *E. coli* in which the half-life of YFP was around 15 minutes [17-20] Additionally, cells displayed large variation in the levels of fluorescence intensity after IPTG induction (Fig 2A and Fig 3B). These features proved to be significant limitations with this tagging system and hence it was not suitable for tracking the fate of pINV in *S. sonnei*.

**Figure 2.**
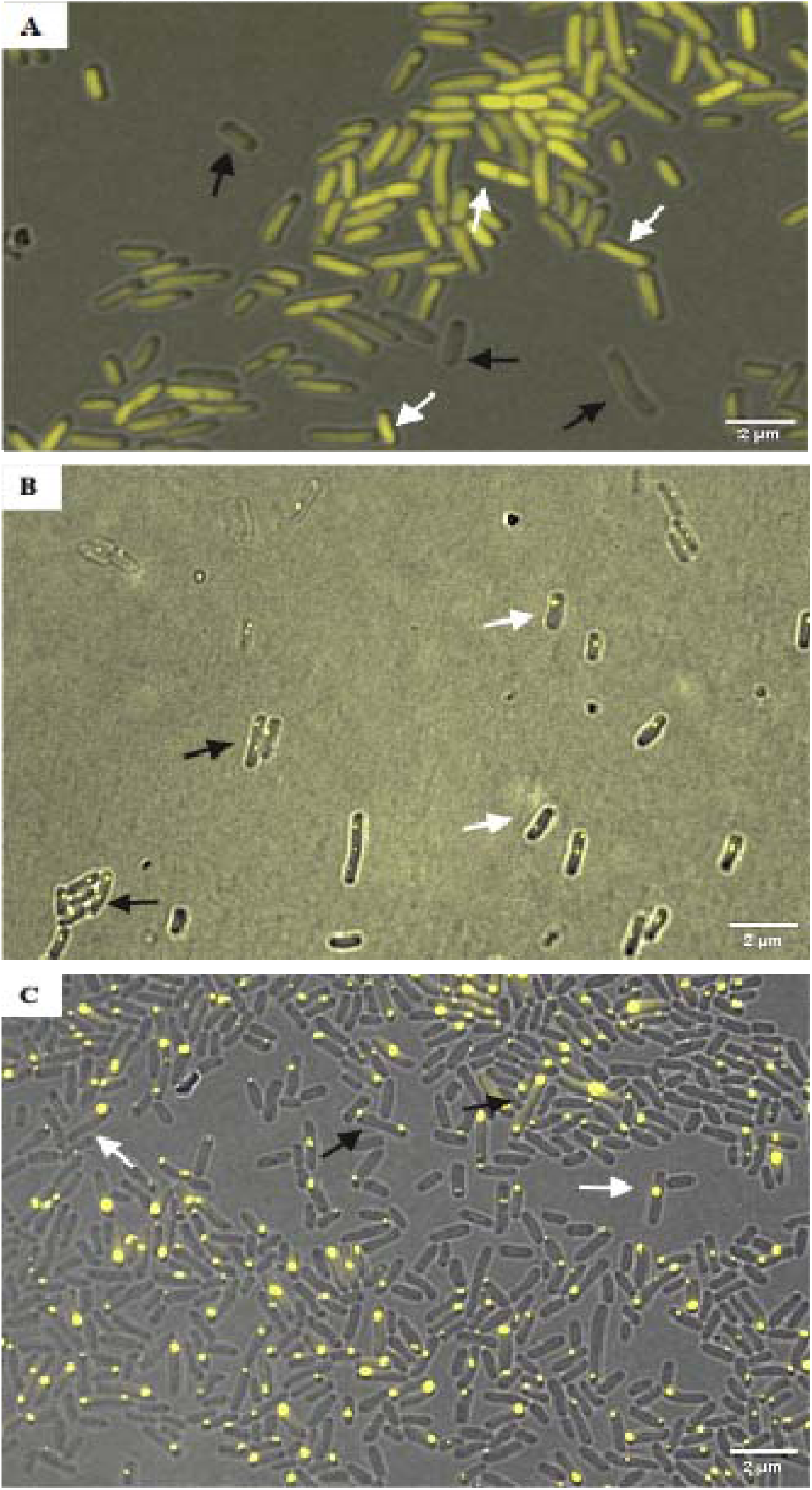
Distinct patterns of YFP-based fluorescent tagging reveal pINV localization in *Shigella sonnei* by live-cell microscopy. Live-cell fluorescence microscopy was used to visualize *S. sonnei* cells carrying pINV tagged with three different YFP-based systems. **(A)** Diffuse cytoplasmic fluorescence is observed in cells expressing an mYPet-ssrA fusion protein encoded directly on pINV. Fluorescence intensity varies across the population: white arrows denote cells with high YFP expression, while black arrows highlight cells with low or heterogeneous expression levels. **(B)** Cells harboring a *tetO*-tagged pINV and co-expressing a TetR-YFP fusion protein (from a second plasmid) exhibit discrete but low-intensity fluorescent foci, typically 1–2 per cell. White arrows indicate cells with a single focus, and black arrows indicate cells with two foci. Increased excitation exposure was used to enhance the weak signal. **(C)** The *parS-*ParB-sfYFP system produces bright, distinct foci in cells containing *parS*-tagged pINV and a ParB-sfYFP fusion expressed from a separate arabinose-inducible plasmid. White arrows point to cells with one fluorescent focus, while black arrows denote cells with two foci.All images were acquired using an EVOS® FL Auto Imaging System with a 100× oil-immersion objective (NA 1.4). YFP fluorescence was captured using 470/22 nm excitation and 525/50 nm emission filters. Scale bars: 2 μm.

**Figure 3:**
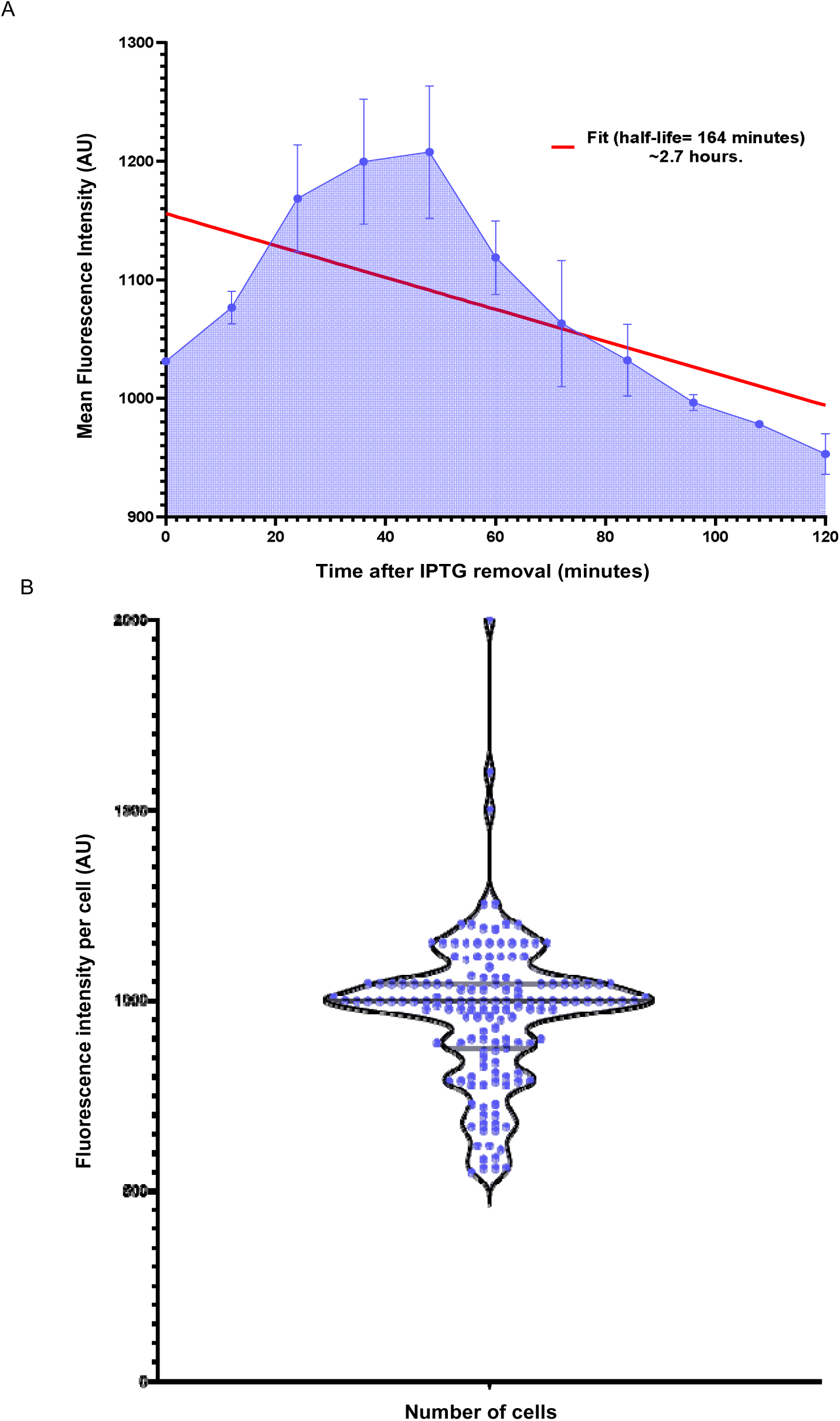
Degradation kinetics and heterogeneity of mYPet-ssrA-tagged pINV fluorescence in *Shigella sonnei*. (A) Mean fluorescence intensity of mYPet-ssrA over time after removal of IPTG inducer. After removal of the inducer IPTG (T=0), live snapshots of *S. sonnei* culture were acquired every 3 minutes using an EVOS® FL Auto Imaging System, 100X (NA1.4, oil-immersion type) lens. Fluorescence images were acquired at excitation wavelength of 470/22 nm and emission wavelength of 525/50 nm for YFP. Image analysis, cell counts, and fluorescence intensity measurements were done using the ImageJ/Fiji platform. Half-life (t_1/2_) was calculated from the graph utilising the formulae: N(t) = N_0_(1/2)^t/t1/2^. Utilizing N_0_: initial quantity at T=0; N_t_: remaining quantity at the end of the experiment. Fluorescence decay follows first-order kinetics with an estimated half-life of ∼164 minutes. Data shown as mean ± SD from 10–50 cells per time point and 3 independent trials. (B) Distribution of single-cell fluorescence intensities of a *S. sonnei* population with a mYPet-ssrA pINV tag. A violin plot representing the fluorescence intensities of 150 cells, with median intensity line overlaid, revealing the marked heterogeneity.

### *tetO*-TetR tagging system did not track pINV for extended periods

We inserted 22 *tetO* sites into pINV downstream of the *vapBC* genes. Cells harbouring the modified plasmid where then additionally transformed with pAB4 plasmid expressing the TetR-YPet fusion protein. Live-cell imaging showed either 1 or 2 YFP foci in most cells, consistent with the known copy number of pINV derived from genome analysis [7]. Although the foci revealed the incidence and location of pINV (Fig 2B), signals were generally of low fluorescence intensity. This could be due to inadequate levels of the TetR-YPet fusion protein in the cell, excessive degradation, or insufficient number of *tetO* binding sites on pINV [21]. Of note, a single *tetO* site will be bound by one TetR dimer hence, there should be at least 2 Ypet molecules per focus [21]. To overcome the intensity issue, higher levels of excitation and exposure were used during image acquisition. However, optimisation of the imaging parameters did not improve signal, but instead caused photobleaching and loss of the YFP signal. Consequently, pINV foci could not be tracked for an extended period (*e.g.* for two hours of live imaging) preventing us from examining its maintenance, movement, and segregation in bacteria.

### The *parS*-ParB tagging system successfully detected pINV

We next inserted two *parS* sites from *C. crescentus* onto pINV of *S. sonnei* and transformed this strain with pSM019 encoding ParB-sfYFP fusion protein under the control of an arabinose inducible promoter. Initially expression of ParB-sfYFP fusion in liquid culture was induced with 0.2% arabinose for 2 hours at 37 °C. During live cell imaging, pINV was detected as 1-2 YFP-foci per cell, representing the copy number of pINV/cell (Figure 2C) [7]. The observed pINV-YFP foci were intense and were robustly detected in every bacterial cell (Figure 2C). The marked fluorescence intensity might have been due to the arabinose inducible promoter present on pSM019 for expressing the ParB-sfYFP fusion protein [22, 29]. Owing to this, photobleaching was not an issue, as the YFP foci were visible even with low exposure of the YFP excitation wavelength. Notably, we found that quality of the imaging data depends on the expression conditions since overexpression of ParB-sfYFP with either a higher concentration of arabinose or induction for more than 2 hours led to an intense but diffused YFP foci across the cell, instead of the distinct 1-2 YFP foci (Fig 4).

**Figure 4.**
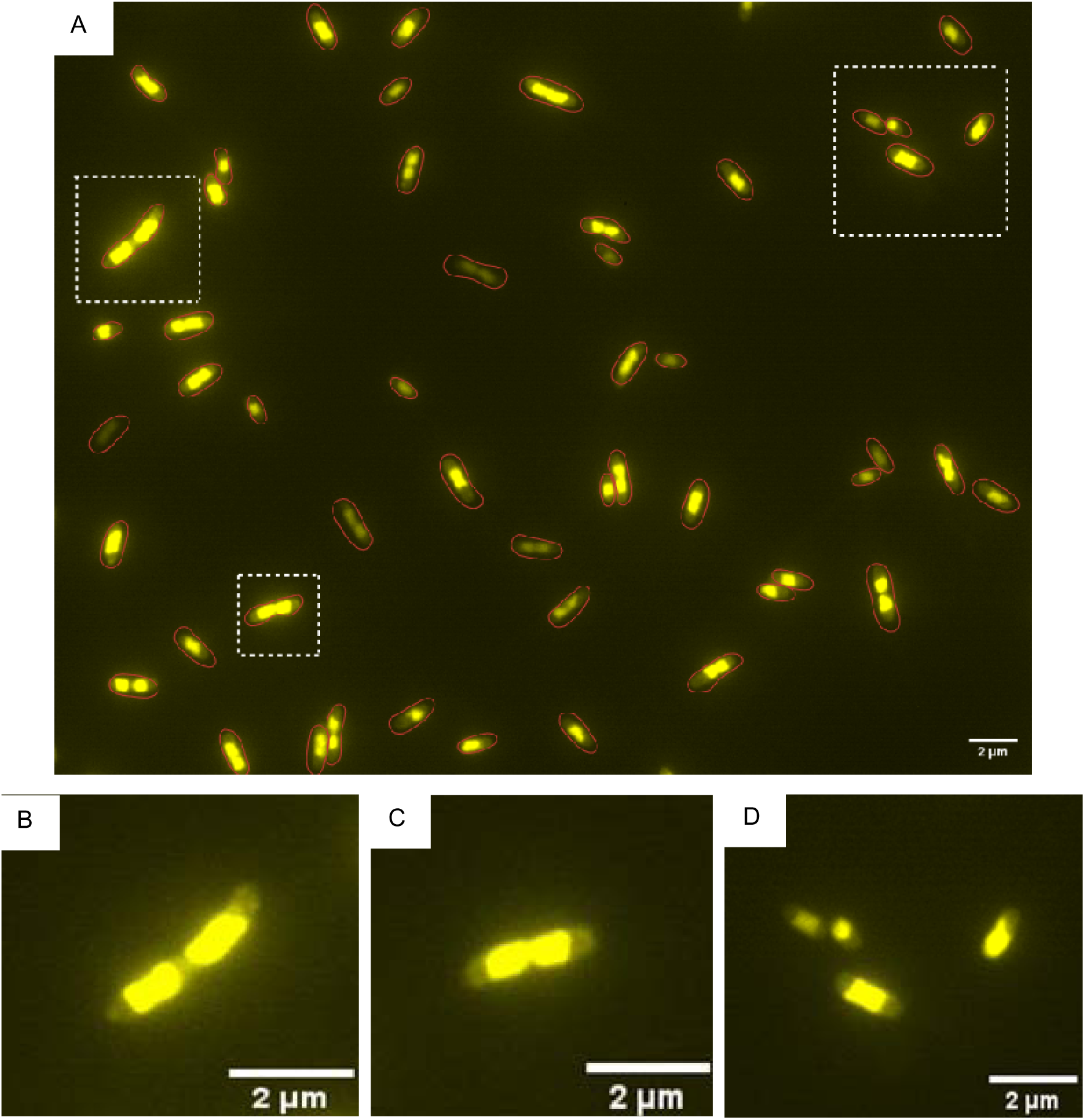
Live-cell fluorescence microscopy of *Shigella sonnei* cells expressing ParB-sfYFP reveals ParB spreading upon over induction. Cells harbouring the pINV-tagged *parS*-ParB-sfYFP system (pSM019) were induced with 0.2% arabinose at 37□°C for 3 hours. The main panel (4A) shows an intense and dispersed YFP fluorescence across the cell, in contrast to discrete foci observed under controlled expression (0.2% arabinose at 37□°C for 2 hours). White dashed boxes highlight representative cells, which are enlarged in panels 4B– 4D. The blurry and extended signal suggests nonspecific spreading of ParB to adjacent chromosomal sites, complicating resolution of the pINV structure. Images were acquired using an EVOS® FL Auto Imaging System with a 100× oil-immersion objective (NA 1.4); excitation/emission: 470/22□nm and 525/50□nm.

These findings suggested that the *parS*-ParB-sfYFP tag was an excellent system for labelling pINV in comparison to the other two methods. *parS*-ParB-sfYFP tag could be employed as an effective tool to track the presence/absence of pINV, its location and movement during live cell imaging.

### Deciphering the movement of pINV across bacterial cell division

We performed live-cell imaging with phase contrast and fluorescence microscopy for *S. sonnei* containing the *parS*-parB-sfYFP pINV tag. Imaging was carried out for 2 hours at 37^0^C, using GMM-1% agarose pads. The acquired images were then analysed with ImageG/Fiji’s additional plugin MicrobeJ and MATLAB based BacStalk software [24-26]. Single cells in the phase contrast images were first distinguished according to their shape, size and lengths, after which pINV-sfYFP foci were detected, and the localized intensities were quantified. This enabled the construction of a demographic view of the bacterial population, depicting the movement and segregation of pINV across the cell cycle of *S. sonnei*. The generated static demograph plot (Fig 5) reveals that pINV is frequently present in a region between the pole and centre of the cell. During cell division, the plasmid migrates to the centre of the cell, where it replicates, then the replicons segregate to the mother and daughter cells. Following cell division, pINV moves back to a pole-proximal region which is toward the old pole in the mother cell, and the newly formed pole of the daughter cell. This pattern is consistent with current literature about DNA segregation of the chromosome and low copy number plasmids which contain a Type I, *parAB* partitioning system [30].

**Figure 5:**
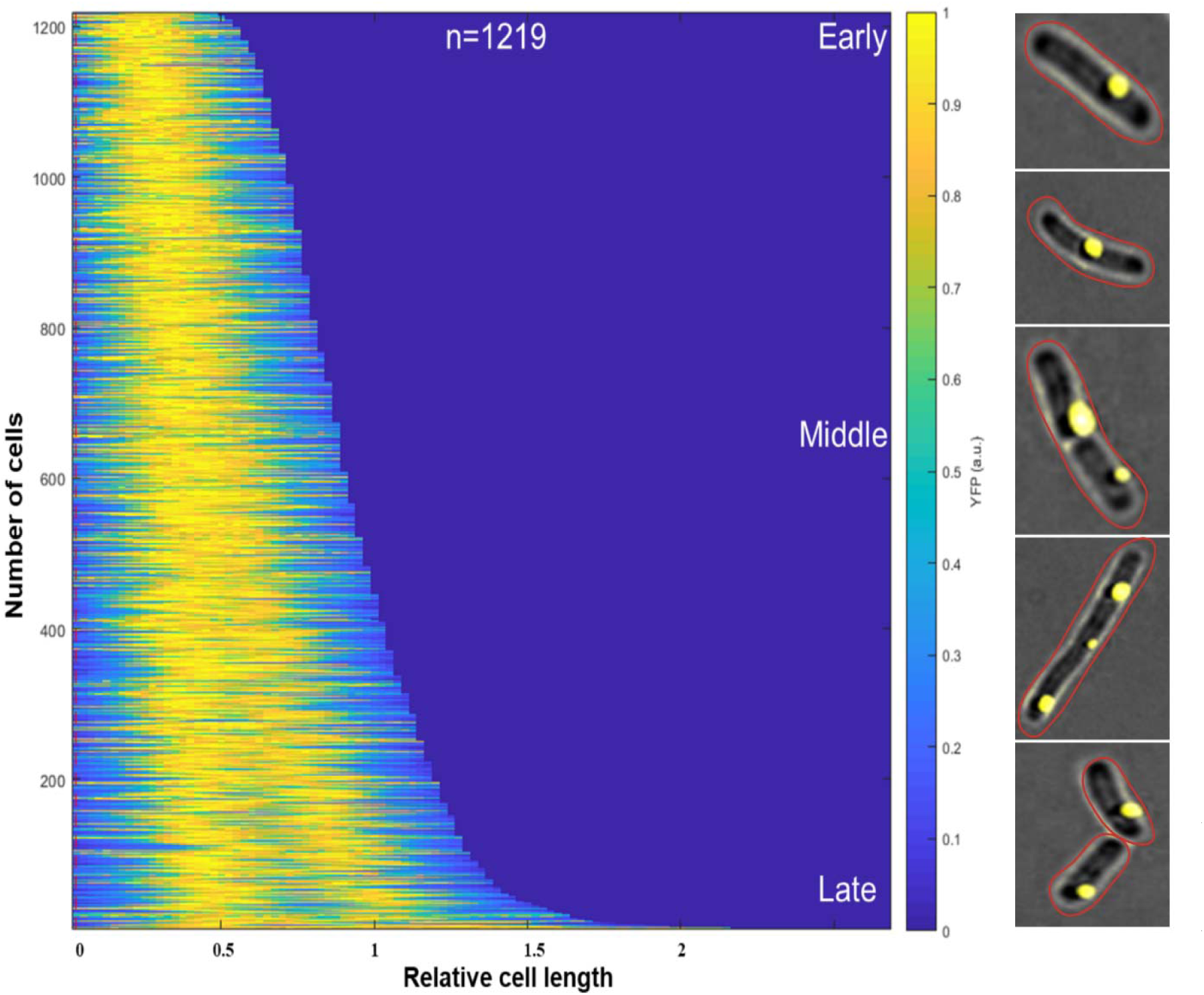
Localization pattern of pINV during bacterial division in *Shigella sonnei*. Microscopic images of a single cell depicting the movement of pINV across the cell during bacterial growth. It is observed that pINV moves from being near the pole to the centre as the cell undergoes division and then is seen back at the pole in both mother and daughter cells. Microscopic images have been acquired using an EVOS® FL Auto Imaging System: 100X (NA1.4, oil-immersion type) lens. Fluorescence images were acquired at excitation 500 nm for YFP plasmid tag and at 585 nm for mKate2. Images were captured from different fields of view and from 3 independent repeats. The demograph showing the normalized fluorescence intensity and the exact location of pINV during bacterial growth. Bacterial segmentation and the demographic analysis exhibiting the fluorescence distribution of pINV foci in a large population of cells was created using BacStalk tool. The cells were sorted by length to highlight the different cell cycle stages, arranged as shorter pre-divisional cells (top) to longer post-divisional cells (bottom) with the red dotted line representing the centre of the cell. Demographic analysis of the YFP foci signal in the imaged cells shows localization of pINV mostly between the cell pole and centre independent of the cell lengths. at 37□°C. Optical

### The *parS*-ParB tag did not impact bacterial growth and plasmid loss

It is critical that introduction of a labelling technique of choice does not interfere with the behaviour of the plasmid during bacterial growth. Hence, we next examined whether the *parS*-ParB tagging system imparted a fitness cost during bacterial growth or affected the maintenance of pINV. Therefore, the bacterial growth rate and plasmid loss rate due to the addition of *parS*-ParB tag were examined and compared with wild-type *S. sonnei*.

We found no significant difference in the growth of the wild-type *S. sonnei* strain and *S. sonnei* strain with *parS*-ParB-sfYFP plasmid tag, when grown in LB or GMM (Fig 6A). This demonstrates that the insertion of *parS* sites on pINV and ParB-sfYFP expression did not cause substantial metabolic burden or toxicity for the bacteria. Additionally, we performed a Congo red (CR) assay to assess the rate of pINV loss by *S. sonnei* strain with the *parS*-ParB-sfYFP plasmid tag, compared to the wild-type strain. In this assay, we measured the emergence of CR-negative colonies to estimate rates of pINV retention/loss after 16 hours. This was quantified by plating bacteria onto CR-TSA, and CR-negative colonies confirmed using multiplex PCR. No significant difference was observed in the rates of emergence for CR-negative colonies between the two strains after around 25 generations (p value: 0.7838, Fig 6B). Furthermore, using multiplex PCR, it was observed that none of the emergent CR-negative colonies had lost the entire pINV plasmid and only had a partial loss of the T3SS genes (data not shown here). These results demonstrated that the insertion of *parS* sites on pINV and ParB-sfYFP expression did not affect maintenance of pINV in *S. sonnei*.

**Figure 6:**
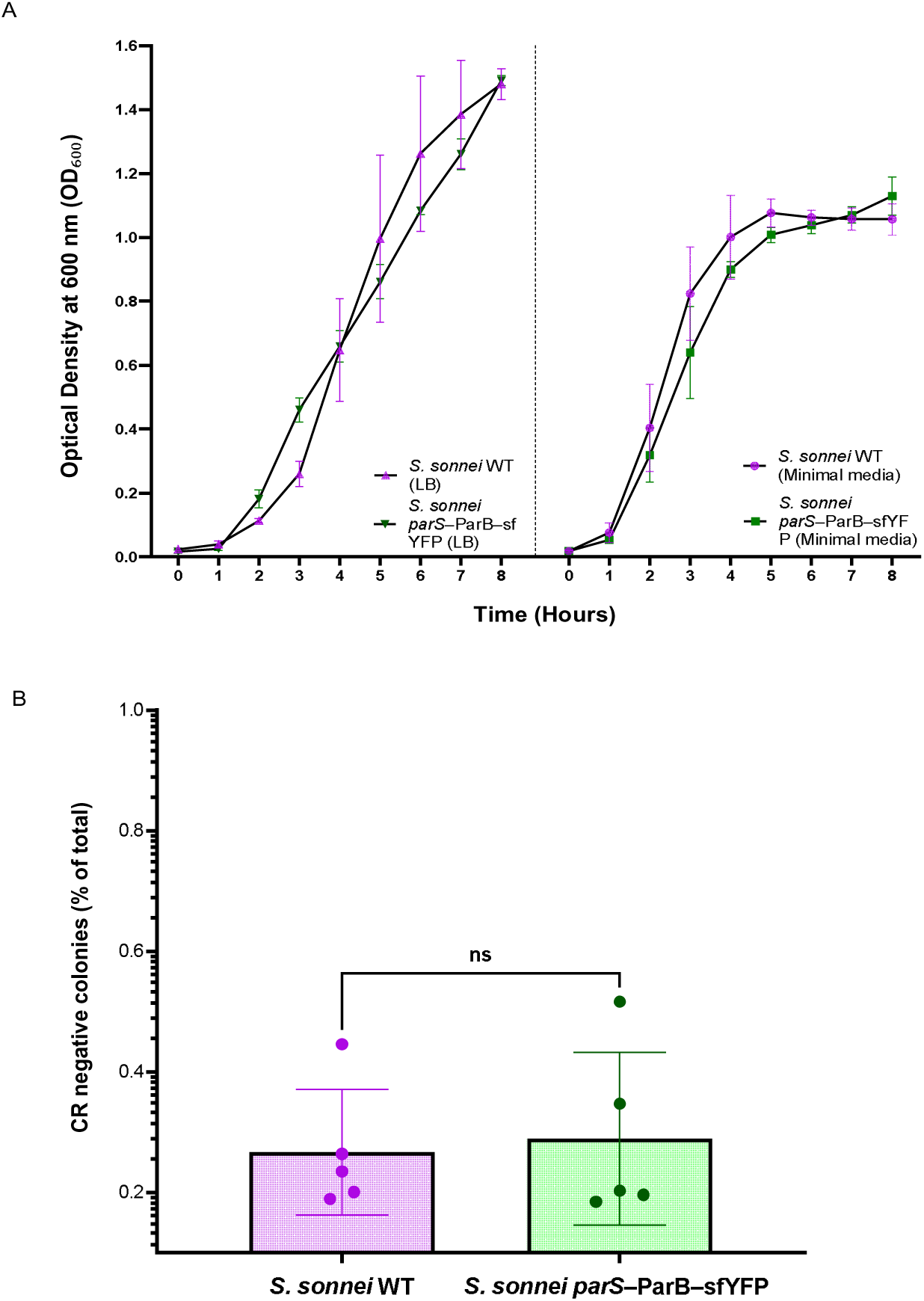
Effect of *parS* site insertion and ParB-sfYFP expression on growth and plasmid stability in *Shigella sonnei*. (A) Growth curves for wild-type *S. sonnei* and *S. sonnei* carrying the *parS*–ParB–sfYFP construct, cultured in LB and glucose minimal media at 37□°C. Optical density at 600 nm (OD□□□) was measured over time. No significant differences in growth were observed between the strains under either condition. Data represent mean ± standard deviation (SD) of three technical replicates. (B) Congo red binding assay evaluating pINV plasmid stability. Percentage of CR□ colonies (indicative of plasmid loss) after ∼25 generations at 37□°C is shown for both strains. Each dot represents an independent biological replicate (n□=□5). Box plots display median ± interquartile range, with whiskers showing full range. Differences were not statistically significant (unpaired Student’s t-test, p□>□0.05).density at 600 nm (OD□□□) was measured over time.

## DISCUSSION

*Shigella* spp. are a public health threat, and it is important to understand its virulence mechanisms to help design strategies to counter the pathogen [3]. *Shigella*’s plasmid pINV harbours virulence genes, which encode the T3SS and many effectors [5, 8]. Several T3SS effectors are also immunogenic and are considered potential vaccine candidates for the prevention of shigellosis [5, 8]. More generally, comprehending the dynamic interaction between bacteria and their plasmids is an essential endeavour, offering knowledge about genetic transfer, adaptation, the spatial/temporal interactions, and microbial evolution. In this study, using an amalgamation of fluorescent tags and live cell fluorescence microscopy, we visualized and tracked pINV in live bacteria. To our knowledge, this is the first description of detection of the large virulence plasmid pINV in *Shigella* spp, at a single-molecule and single-cell level. This study enabled us in tracking pINV such as its presence and loss, further aiding to characterise the real-time cellular dynamics of the plasmid.

We investigated three tagging systems to either detect cells harbouring pINV or directly label the plasmid. Each tagging systems had its own advantages and limitations, that could be tailored according to the needs of a specific research question. In this instance, it was found that the *parS*-ParB-sfYFP pINV tag accurately detected the presence, location, and segregation of pINV in live bacterial cells.

The first tagging approach employed labelling cells with pINV with a fast-degrading fluorescent protein. The addition of a ssrA tag the C-terminus of a protein leads to its rapid degradation by cellular proteases [18-20]. In *E. coli,* ClpXP is a proteolytic complex that recognizes and degrades >90[% of ssrA-tagged substrates [18-20]. In this study, we fused a ssrA tag to mYPet fluorescent protein expressed on pINV and used this modified system to then visualise live *S. sonnei* cells containing the plasmid. This system generated dispersed YFP fluorescence across the cell, so only indicates the presence/absence of pINV and not its cellular location. Contrary to expectations, the rapid-degradation tag was not able to promptly detect the loss of pINV as decay of its signal only occurred over hours after the IPTG inducer was removed. The YFP intensity was also highly heterogeneous across cells which could be due to the variable expression of either mYPet-ssrA or proteases for degrading the fluorescent protein [17-20]. In the end, due to these technical issues, the mYPet-ssrA tag was not used further to characterise the cellular dynamics of pINV. Further development of this system could focus on improving the specificity of proteolysisby additional regulatory protein factors such as SspB [32]. This protein binds to ssrA-tagged substrates and augments recruitment of ClpX, enhancing proteolysis [32].

The *tetO*-TetR-YPet pINV tag was able to reveal the location and copy number of pINV inside cells but was not suitable for tracking the movement of the plasmid during the cell cycle. This was due to the low intensity (low signal-to-noise ratio) of the YFP foci, requiring high levels of fluorescence exposure time for visualization, which in turn cause rapid photobleaching and loss of signal. Of note, each *tetO* operator sequence binds to a single TetR-YPet dimer [21]. Hence, to boost this current system and achieve a high signal-to-noise ratio, increasing the number of binding sites for TetR-YPet, using more than 22 repeats could be considered. However, this could be detrimental, as integration of a large DNA repeat sequence could cause instability of plasmid and hinders its widespread application as a DNA imaging tag [33]. Furthermore, adding an inducible promoter in the TetR-YPet plasmid (pAB4) to express higher concentrations of the fluorescent fusion protein could also be carried out [21]. Exploiting both these strategies has the potential to improve the intensity of the YFP foci and aid in enhanced detection of pINV in the future [21, 33].

A heterologous type I *parS*-ParB segregation system from *C. crescentus* was used in this study to visualise and track pINV in *S. sonnei* [22]. ParB proteins specifically bind to centromere-like DNA sequences called *parS* sites, polymerises onto the DNA and are involved in the segregation of bacterial plasmids and chromosomes [29, 30]. Thus *parS*-ParB-based anchor system can be used in live-cell DNA imaging, assisting in the labelling of DNA efficiently with negligible disturbance to genomic stability [29, 30]. In this study, the insertion of 90 bp region comprising of two *parS* sites onto pINV, allowed the arabinose-induced ParB-sfYFP to bind around the *parS* sites for loci-specific visualization. It permitted detecting the presence/absence of pINV and tracking it over 2 hours of live-cell imaging with an appropriate fluorescence intensity. [16]. As explained before, the *parS* sites on pINV were inserted downstream of the vapBC (TA) genes, along with an antibiotic cassette ensuring that the inserted sequences are stably maintained on pINV. TA systems such as VapBC are primarily known to be transcriptionally regulated [11]; hence inserting *parS* sequences near these genes and the corresponding ParB binding and DNA polymerisation could cause an effect on bacterial growth and plasmid retention. This was ruled out based on the growth curve analysis and CR-plasmid loss measurements.

It has been shown that apart from exclusive binding to the *parS* sites, ParB proteins have the ability to associate non-specifically with DNA adjacent to *parS* sites, a phenomenon called spreading [34]. In this study, the pSM019 expressing the ParB-sfYFP fusion protein under an arabinose promoter contains the p15A/pACYC origin of replication and is a low copy number plasmid (15 copies per cell) [22]. The phenomenon of spreading was indeed encountered in our case, as an overproduction of the ParB-sfYFP led to a diffused YFP foci and its associated fluorescence instead of a single focus representing pINV, which would indicate spreading of ParB across other DNA loci or adjacent sites. This hindered locating and tracking the plasmid organisation across the cell. In this study, the *parS*-ParB tagging system proved to be the most suitable choice for visualizing and examining the pINV parameters. It facilitated acquiring information about pINV’s location (sub-proximal pole region), copy number (1-2/cell), segregation (tracking during cell division), and maintenance (presence/loss) in live bacterial cells. This tagging system has additionally been applied to visualize gene transcription or viral replication in mammalian cells [29], genome organization in drosophila or mammalian cells [29], and additionally been used to study DNA transfer via plasmid conjugation [31].

It is known that for any single copy plasmid such as pINV, its replication must be matched with the division of the chromosome under an active partitioning system [35. 36]. A tripartite partitioning system is present on pINV, which is a conserved molecular mechanism that ensures a faithful plasmid segregation into daughter cells. [37-40]. This system consists of ParA (motor protein encoding ATPase or GTPase), ParB (centromere-binding adaptor protein) and a *cis*-acting site *parS* (centromere-like sequence area) [37-40]. Take in note that this partitioning ParABs system is sufficiently different from the one used for tagging pINV in this study. We utilized *Caulobacter*’s parS-ParB sequences to tag pINV and as seen in this study, there was no interference or cross communication that occurs between the two systems.

The most well-studied bacterial plasmid segregation system, with respect to mechanism, is that harboured by the R1 plasmid [41]. The plasmid shows a dynamic pattern of localization within the *E. coli* cell, being positioned near the cell pole. The plasmid then moves to the central position during cell growth (for duplication by the replisome), and the daughter plasmids move back to their pole-proximal location [41]. In another study, the dynamics of ParA from F plasmid in *E. coli* was examined using fluorescence microscopy and computational modelling [42]. They illustrated that ParA ensures partition of plasmids takes place before the full segregation of bacterial chromosomes at or prior to cell division [42].

With the means of the generated demographic plot in this study, it was observed that pINV in *Shigella*, is commonly present in the sub-proximal pole of the cell. During cell division, pINV moves to the centre of the cell, where the plasmid replication and segregation takes place before the cell division. Following the division, pINV again moves back to near the bacterial poles in both the mother and daughter cell. This demographic view of the *S. sonnei* population exhibited a similar pattern of pINV dynamics as shown in earlier studies where segregation carried out by the ParABS system has been investigated during cell division [40-42].

## CONCLUSION

In conclusion, this study visualizes and aids in understanding the real-time cellular dynamics of the large virulence plasmid in *Shigella* using fluorescent plasmid tags and live-cell microscopy. While we have focused on methods for tracking pINV across cell division, these techniques can be utilized in investigating cellular processes, disease mechanisms, and therapeutic interventions associated with other bacterial plasmids.

## Author contributions

TS: Conceptualization, Data curation, Formal analysis, Investigation, Methodology, Project administration, Validation, Writing – original draft, Writing – review & editing

VL: Data curation, Formal analysis, Methodology, Writing – review & editing SU: Resources, Supervision, Writing – review & editing

CT: Conceptualization, Funding acquisition, Project administration, Supervision, Writing – review & editing

