## Supplementary Table 1 for "Real-time visualisation of the intracellular dynamics of *Shigella’s* virulence plasmid"

**Supplementary Table 1: Bacterial strains and plasmids used in this study**

| Strain No | Species | Strain | Plasmid | Insert | Resistance marker | Description | Reference |
| --- | --- | --- | --- | --- | --- | --- | --- |
| 1 | *S. sonnei* | CS14 | - | - | - | Wild-type *Shigella* strain | 14 |
| 2 | *E. coli* | DH5alpha | pKD46 | - | Ampicillin | Helper plasmid expressing the  lambda red genes (gama, beta, and  exo) | 15 |
| 3 | *E. coli* | BW25141 | - | - | - | Wild type *E. coli* strain with R6K gamma replication origin (needed to support pKD3/4 ori. | 15 |
| 4 | *E. coli* | BW25141 | pKD4 | - | Kanamycin | Used for template generation for  homologous recombination | 15 |
| 5 | *E. coli* | BW25141 | pKD3 | - | Chloramphenicol | Used for template generation for  homologous recombination | 14 |
| 6 | *E. coli* | AB1157 | - | - | Gentamycin | pLAU44 derivative *E. coli* strain containing multiple tetO repeats | 20 |
| 7 | *E. coli* | DH5alpha | pAB4 |  | Chloramphenicol | TetR-YPet encoding plasmid | 20 |
| 8 | *E. coli* | BW25141 | pKD3 | 22xtetO repeats inserted into pKD3 | Kanamycin | Used for knock-in template generation | This study |
| 9 | *S. sonnei* | CS14 | pAB4 | 22xTeto repeats inserted into pINV | Kanamycin and Chloramphenicol | *Shigella* strain with the tetO-TetR-YPet pINV tag | This study |
| 10 | *E. coli* | DH5alpha | pROD50 | - | Ampicillin and Kanamycin | Inducible plasmid encoding mYPet | 17 |
| 11 | *E. coli* | DH5alpha | pROD50 | 11 amino acid peptide-AANDENYALAA fused with mYPet | Ampicillin and Kanamycin | mYPet fused with the ssrA tag | This study |
| 12 | *S. sonnei* | CS14 | - | mYPet-ssrA tag into the pINV. | Kanamycin | *Shigella* strain with the fast degrading mYPet-ssrA pINV tag | This study |
| 13 | *E. coli* | DH5alpha | pSM040 | - | Chloramphenicol | Plasmid containing parS sites | 21 |
| 14 | *E. coli* | DH5alpha | pSM019 | - | Ampicillin | Inducible plasmid encoding ParB-sfYFP | 21 |
| 15 | *E. coli* | BW25141 | pKD3 | 90kb region containing 2 parS sites inserted into pKD3 | Kanamycin | Used for knock-in template generation | This study |
| 16 | *S. sonnei* | CS14 | pSM019 | 90kb region containing 2 parS sites inserted into pINV | Ampicillin and Kanamycin | *Shigella* strain with the parS-ParB-sfYFp pINV tag | This study |
