## Supplementary Table 2 for "Real-time visualisation of the intracellular dynamics of *Shigella’s* virulence plasmid"

**Supplementary Table 2: Primers used in this study.**

| **Primer Name** | **Sequence (5’-3’)** | **Role** |
| --- | --- | --- |
| tetO repeats-F | aactaaggaggatattcatatggaccatggGTGATAGAGAAGACGAACCG | Amplifying the 22xtetO repeats |
| tetO repeats-R | ggaacacttaacggctgacatgggaattagTCTGCTAGAGTCTCTCTCCA | Amplifying the 22xtetO repeats |
| pKD4-tetO-F | TGGAGAGAGACTCTAGCAGActaattcccatgtcagccgt | Gibson primers to insert 22xtetO repeats into pKD4 |
| pKD4-tetO-R | CGGTTCGTCTTCTCTATCACccatggtccatatgaatatc | Gibson primers to insert 22xtetO repeats into pKD4 |
| 22tetO-knockin-F | caatgtgcaccgggaacgcggggaggatgttgagccgggagatgatttctgatgaacagggtgtaggctggagctgcttc | Primer to amplify the knockin fragment with upstream homologous sequence from pINV |
| 22tetO-knockin-R | atgggaattagTCTGCTAGAGataacacccgggaatttgaacgtgtgggcggcctgagaactgaagactggagctgacctg | Primer to amplify the knockin fragment with downstream homologous sequence from pINV |
| tetO insertion check-F | agccgggagatgatttctga | Primer to confirm correct insertion and position of 22xtetO repeats into pINV |
| tetO insertion check-R | GGAAACCACCGTATTTCTCA | Primer to confirm correct insertion and position of 22xtetO repeats into pINV |
| ssrA-mYPet-F | GCAGCCAACGATGAAAATTATGCACTGGCAGCATAACCCGGGTGTAGGCTGGA | Inserting ssrA peptide into pROD50 plasmid |
| ssrA-mYPet-R | TGCTGCCAGTGCATAATTTTCATCGTTGGCTGCtttgtacaattcatTcatac | Inserting ssrA peptide into pROD50 plasmid |
| mYPet-ssrA insert-pINV-F | caatgtgcaccgggaacgcggggaggatgttgagccgggagatgatttctgatgaacaggACGCAATTAATGTGAGTTAG | Primer to amplify the knockin fragment with upstream homologous sequence from pINV |
| mYPet-ssrA insert-pINV-R | ataacacccgggaatttgaacgtgtgggcggcctgagaactgaagactggagctgacctgTCCTCCTTAGTTCCTATTCC | Primer to amplify the knockin fragment with downstream homologous sequence from pINV |
| mYPet-ssrA insertion check-F | agccgggagatgatttctga | Primer to confirm correct insertion and position of mYPet-ssrA into pINV |
| mYPet-ssrA insertion check-R | GGAAACCACCGTATTTCTCA | Primer to confirm correct insertion and position of mYPet-ssrA into pINV |
| parS sites-F | actcatcggcgtttcacgtg | Amplifying the 90kb region containing 2 parS sites |
| parS sites-R | agcagctccagcctacacaccaagtgatgtttcacgtgga | Amplifying the 90kb region containing 2 parS sites |
| pKD4-parS-F | tccacgtgaaacatcacttggtgtgtaggctggagctgcttcg | Gibson primers to insert 90kb region containing 2 parS sites into pKD4 |
| pKD4-parS-R | tcctccttagttcctattcc | Gibson primers to insert 90kb region containing 2 parS sites into pKD4 |
| parS-knockin-F | caatgtgcaccgggaacgcggggaggatgttgagccgggagatgatttctgatgaacaggactcatcggcgtttcacgtg | Primer to amplify the knockin fragment with upstream homologous sequence from pINV |
| parS-knockin-R | tgtttcacgtggaacatcccataacacccgggaatttgaacgtgtgggcggcctgagaactgaagactggagctgacctg | Primer to amplify the knockin fragment with downstream homologous sequence from pINV |
| parS-insertion check-F | agccgggagatgatttctga | Primer to confirm correct insertion and position of the 90kb region containing 2 parS sites into pINV |
| parS-insertion check-R | GGAAACCACCGTATTTCTCA | Primer to confirm correct insertion and position of the 90kb region containing 2 parS sites into pINV |
| *ori*-F | GTGACCTCCTCAGAATAATCC | Multiplex PCR for confirming pINV loss |
| *ori*-R | AAAAGATACATTGCACCCTGT | Multiplex PCR for confirming pINV loss |
| *hns*-F | GCTCAACAGTATGCACAGAA | Multiplex PCR for confirming pINV loss |
| *hns*-R | TTGCAAAGGCGTTGAATTA | Multiplex PCR for confirming pINV loss |
| *virB*-F | ACATCAGAGCTCCACAAGAA | Multiplex PCR for confirming pINV loss |
| *virB*-R | AGACGATAGATGGCGACGAAA | Multiplex PCR for confirming pINV loss |
| *virF*-F | CTTAGCTTGTTGCACAGAGA | Multiplex PCR for confirming pINV loss |
| *virF*-R | AAGARGGGCTTGATATTCCG | Multiplex PCR for confirming pINV loss |
| *gmvAT-*F | CGGTAAAGCCTGAATGGAAA | Multiplex PCR for confirming pINV loss |
| *gmvAT-*R | TAACAGCCTGAATGGAAA | Multiplex PCR for confirming pINV loss |
